# Experimental evidence for sexual selection against inbred males when it truly counts

**DOI:** 10.1101/045716

**Authors:** Regina Vega-Trejo, Megan L. Head, J. Scott Keogh, Michael D. Jennions

**Author notes:** Corresponding author; Regina Vega-Trejo. Data Accessibility: Data will be deposited in Dryad upon acceptance.

## Abstract

Although there are many correlational studies, unbiased estimates of inbreeding depression only come from experimental studies that create inbred and outbred individuals. Few such studies determine the extent to which inbreeding depression in males is due to natural or sexual selection. Importantly, traits that are closely related to fitness are predicted to be most strongly affected by inbreeding depression, so measuring fitness or key fitness components, rather than phenotypic traits, is necessary to estimate inbreeding depression accurately. Here, we experimentally created inbred and outbred male mosquitofish (*Gambusia holbrooki*) by mating full-sibs (*f*=0.25). We show this led to a 23% reduction in genome-wide heterozygosity. Males were then raised on different diets early in life. We then allowed adult males to compete freely for females to test if inbreeding, early diet, and their interaction affect a male’s share of paternity. Early diet had no effect on paternity, but outbred males sired almost twice as many offspring as inbred males. We also found that males with a relatively long gonopodium (intromittent organ) had greater reproductive success. We demonstrate that inbreeding has important consequences because it negatively affects a key component of male fitness. Given there was no difference in adult mortality this finding can only be due to inbreeding negatively affecting sexually selected traits.

## Introduction

Evidence that inbreeding in animals negatively affects lifetime reproductive success or other close proxies of fitness, comes from two main streams of research: correlational studies (mainly of wild populations), and experimental studies on laboratory or captive populations that create inbred and outbred individuals. Correlational evidence for inbreeding depression includes: a) comparing traits among populations of the same species, specifically between small, isolated populations that have experienced varying degrees of inbreeding and large, outbred populations [1–4]; b) using molecular markers to obtain direct estimates of individual levels of inbreeding within a population and then relating these to fitness measures [i.e. heterozygosity-fitness correlations: HFCs; 5, 6–8]; c) calculating an individual’s inbreeding coefficient from pedigree data and then relating this to a fitness measure [8–11]. Although correlational studies often suggest that inbreeding lowers fitness, other factors cannot be ruled out. For example, inbred individuals might more often occur in peripheral environments that are of low quality such that there is a direct environmental effect on offspring phenotypes. More generally, the reduced fitness of inbred individuals might partly result from additive genetic effects rather than non-additive interactions within loci [i.e. lower heterozygosity; 12]. If focal traits are heritable, and individuals with lower values tend to mate with relatives because they have poorer dispersal ability or struggle to attract mates, this will lead to systematic overestimation of the negative effects of inbreeding [see the discussion in 12]. Inbred offspring will inherit lower trait values regardless of any direct effects of inbreeding. Studies that experimentally manipulate levels of inbreeding with controlled breeding designs offer a better approach when trying to quantify the costs of inbreeding.

To date, relatively few experimental studies have looked at the effects of inbreeding on estimates of fitness in non-domesticated animals. Of these, only a handful of studies have specifically looked at male fitness [e.g. 13, 14–16]. It therefore remains an open question as to the extent to which males are more susceptible than females to inbreeding depression. Mating success and fertilization success under sperm competition are major determinants of male fitness in most species [17–19]. Sexually selected traits that confer a mating or fertilization advantage are often under strong directional sexual selection and, in addition, they tend to be condition-dependent. Condition-dependence has been described as a form of ‘genic capture’ because condition reflects how well the individual accumulates resources [20, 21]. This ability is likely to depend on many traits (e.g. foraging ability, food absorption efficiency, timing of development) all of which could be negative affected by inbreeding. It is therefore plausible that due to sexual selection male mating success will show greater inbreeding depression than an equivalent naturally selected female trait such as fecundity. These data cannot, however, be obtained from studies that measure male lifetime reproduction output as they confound lifespan with reproductive success per potential breeding event (i.e. sexual selection).

Within experimental studies of animals that try to measure fitness there is high variation in the reported magnitude of inbreeding depression [e.g. 16, 22, 23, 24]. One possible source of variation is whether test individuals are exposed to a stressful environment [25, 26]. Inbreeding might make individuals less effective at buffering themselves against environmental stress [27]. Dietary and temperature stress, for example, increase the extent of inbreeding depression in some species [28–31]. More generally, rearing animals in a benign lab environment (or plants in well-watered greenhouses) is often put forward to explain the absence of inbreeding depression in a laboratory study [32, 33]. Another potential source of variation in estimates of inbreeding depression might arise from the evolutionary history of study populations affecting the baseline level and variability of homozygosity. For instance, as mean homozygosity in a population increases the difference in homozygosity between offspring of closely related individuals and those from random matings decreases [34]. This makes it harder to detect inbreeding depression. To date, studies that investigate how these potential sources of variation influence the effects of inbreeding on fitness-enhancing traits remain scant [but see 28, 34, 35].

Here we conduct an experiment to investigate how differences in inbreeding and juvenile diet (i.e. early stressful environment) influence a key component of male fitness, namely their reproductive success. We experimentally generated inbred and outbred male mosquitofish (*Gambusia holbrooki*) that were then reared on different diets as juveniles [36]. We then allowed males to compete freely for access to females and examined their share of paternity. The ability to gain paternity under sperm and mating competition is a key fitness component for males in species with high levels of female polyandry, such as *G. holbrooki*. Importantly our experimental design allows us to isolate sexual selection (as opposed to other forms of natural selection) as the cause of any inbreeding depression. In addition to our experimental manipulation of inbreeding using a controlled pedigree we measured each male’s actual genome wide heterozygosity (based on >3000 SNPS) to shed light on how much variation in inbreeding is needed to detect inbreeding depression. We predict that under the competitive mating scenario we created that, if it occurred, inbreeding depression would be greater for males reared in a stressful environment.

## Methods

### Origin of fish

We used mosquitofish descended from wild caught fish collected in Canberra, Australia. The design used to create inbred and outbred males that were then reared on different diets is fully described in [36]. In brief, in each experimental block we mated individuals from two full sibling families (e.g. A and B in block 1, C and D in block 2 and so on). Brothers and sisters from full sibling families were paired to create inbred offspring (AA, BB; *f* =0.25) and outbred offspring with reciprocal male-female crosses (AB, BA) to generate four cross-types. We set up 29 blocks (= maximum of 116 different family pairings types). The 452 male offspring from 192 broods (some experimental blocks had more than one pairing of a given type) were then reared individually in 1L tanks until maturity. Males then underwent a diet manipulation for 21 days between day 7 and day 28-post birth that lead to almost zero growth [36]. Fish on the control diet were fed *ad libitum* with *Artemia* nauplii twice a day (i.e. standard laboratory feeding regime) while fish on the restricted diet were fed 3mg of *A. nauplii* once every other day (i.e. < 25% of the control food intake). Broods were split evenly between the control and restricted diet treatment.

### Experimental design

To determine whether inbreeding, diet, or their interaction predict paternity we set up mating trials in which four males, one per treatment, could compete and mate freely with a female in a 60L tank (n=31). Males were randomly assigned to each replicate and were not match for size (size range: 18.51 - 26.96 mm). Males were allowed to mate freely with a female for a week and then given a week to recover after the female was removed. The process was then repeated with two more females. The 93 test females were then placed in individual 1L tanks and allowed six weeks to give birth. They were checked for offspring twice daily. Once fish were removed from the treatment they were euthanized and preserved in absolute ethanol and stored at ‐20°C.

### Male morphology

The phenotype of all males was measured prior to being placed in tanks with females. Males were anaesthetized by submersion in ice-cold water for a few seconds to reduce movement and then placed on polystyrene with a microscopic ruler (0.1 mm gradation) and photographed. We measured male standard length (SL = snout tip to base of caudal fin) and gonopodium length (intromittent organ modified from the anal fin) using Image J software [37]. The males were 28 – 37 weeks post-maturity and were marked with a small coloured dot for visual identification using fluorescent elastomer (Northwest Marine Technology, WA) injected subcutaneously behind the caudal fin. They were given at least four days recovery before going into 60L tanks to start mating trials. We calculated relative gonopodium size as the residuals from a linear regression of gonopodium size (log) on male standard length (log).

### Paternity analysis

To determine male reproductive success and heterozygosity for the fish in our experiment we took tissue samples from each male (n=121), females that bred (n=79 of 93), and a maximum of 10 fry per female (n=628 offspring). Two of the 124 males (both outbred) were not found in the tank at the end of the trial (i.e. escaped or died) and therefore no tissue was available. For adults, DNA was extracted from the tail muscle/caudal fin. For fry DNA was extracted from the whole body, excluding the head. DNA was extracted using Qiagen DNeasy Blood & Tissue Kits following the manufacturer’s instructions. After extraction, DNA samples were SNP genotyped. Full methods for the paternity analysis are in [38].

### Heterozygosity

We estimated heterozygosity by using the number of markers that were scored as heterozygous divided by the total number of successfully classified markers for each fish. Based on over 3000 SNP loci we found that a brother-sister mating led to a significant decline in offspring heterozygosity (F_(1,120)_ = 215.1, *P*<0.001). The mean heterozygosity of inbred fish was 23.2% less than that of outbred fish (close to the expected 25% decline). The proportion of loci that were heterozygous was 0.239 ± 0.003 in inbred males and 0.311 ± 0.004 in outbred males (n= 62, 59).

### Statistical analysis

We used Generalized Linear Mixed-effect models (GLMM) with Poisson error to test for fixed effects of inbreeding, relative heterozygosity (see below), diet, body size, relative gonopodium length, and the interaction between inbreeding and diet on how many offspring males sired. We used the *glmer* function in the *lme4* package in *R 3.0.2* software [39]. To obtain a measure of relative heterozygosity we centered heterozygosity (mean = 0) within each inbreeding treatment. We could then test whether it explained additional variation in male reproductive success beyond that associated with the decline in absolute heterozygosity due to inbreeding. We also included the interaction between standardized heterozygosity and inbreeding to test for any difference in the effects of this additional variation in heterozygosity between inbred and outbred males (i.e. the effect will differ if there is a non-linear relationship between absolute heterozygosity and fitness). To account for overdispersion we included individual as a random effect [40]. We included tank as a random effect to account for potential non-independence. We included sire and dam as random effects. There was no effect so we present the simplified version of the model. All model terms were tested for significance using the Anova function in the *car* package specifying Type III Wald chi-square tests. We removed non-significant interactions following [41]. All tests are two-tailed and alpha is set at 0.05.

## Results

### Inbreeding

On average, outbred males sired significantly more offspring than inbred males (Table 1, Fig. 1). In 20 of 31 trials, the two outbred males sired more offspring than the two inbred males. More heterozygous males therefore had significantly greater reproductive success.

**Table 1.**
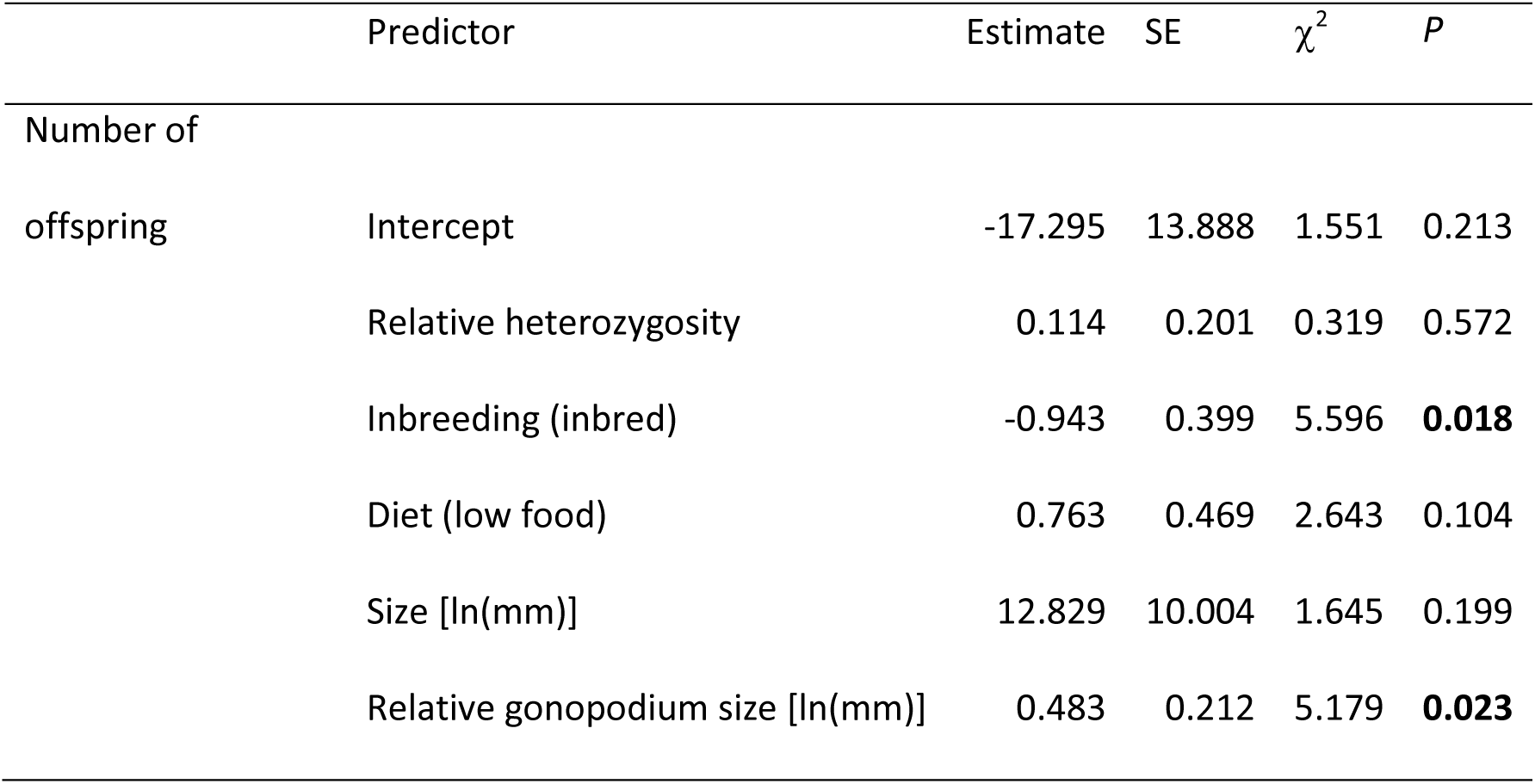
Results from the mixed model with parameter estimates and chi square (χ^2^) tests for heterozygosity, inbreeding, food treatment, size, and relative gonopodium size. P-values in bold indicate significant values.

**Figure 1.**
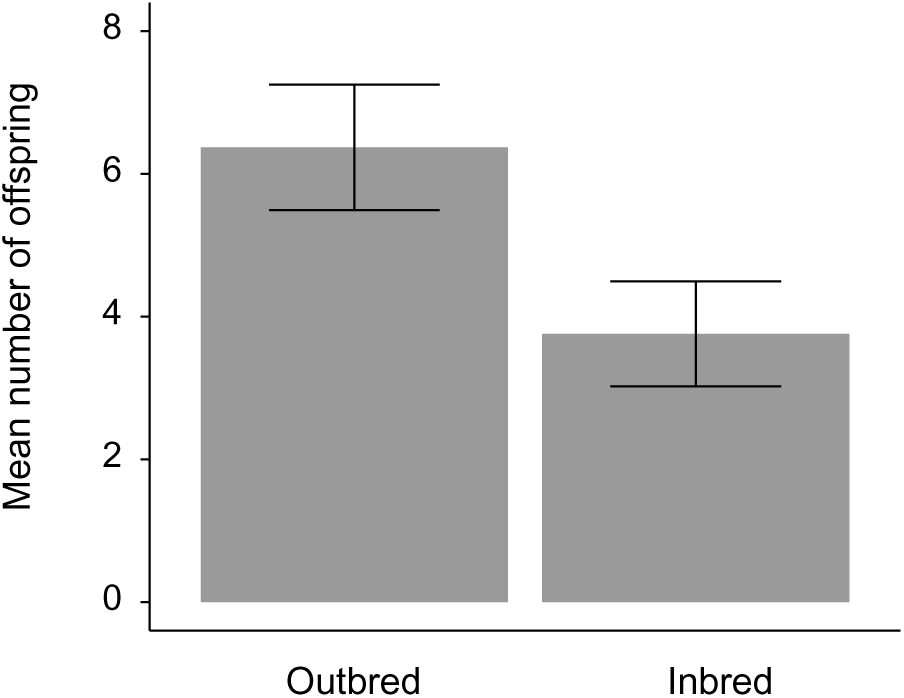
Mean number of offspring (± SE) sired by outbred and inbred males.

### Relative Heterozygosity

We did not find any difference in how relative heterozygosity affected male reproductive success between inbred and outbred males (heterozygosity × inbreeding, *P* = 0.350). There was also no effect of relative heterozygosity on male reproductive success (Table 1). Together these findings indicate that the standing variation in heterozygosity (i.e. that in outbred males) did not predict variation in male reproductive success.

### Diet

We did not find an effect of diet on the number of offspring sired (Table 1). There was also no significant interaction between inbreeding status and diet (*P* = 0.586).

### Male morphology

Males with a relatively longer gonopodium sired significantly more offspring. We did not, however, find an effect of male body size on the number of offspring sired (Table 1).

## Discussion

Inbreeding is expected to decrease fitness due to the negative effects of lower heterozygosity [42, 43]. Here we used a controlled breeding design combined with a genome wide SNP-based measure of heterozygosity to test whether inbreeding, as well as residual variation in heterozygosity, affects a key component of male fitness, namely male reproductive success. We found that one generation of inbreeding between full-siblings (*f* =0.25) significantly lowered a male’s ability to gain paternity by almost 50% (6.37 vs 3.76 offspring). Outbred males sired significantly more offspring than inbred males when they had to compete for mates and fertilization opportunities. Sexual selection therefore favours outbred males. In addition, relative gonopodium length explained some of the remaining variation in reproductive success. Males with a longer gonopodium were significantly more successful. We found no evidence for an effect of diet or body size on male reproductive success. Nor did we find any effect of residual variation in heterozygosity once we accounted for the 23.2% decline in heterozygosity associated with inbreeding in our pedigree design (i.e. full-sibling parents versus unrelated parents).

### Heterozygosity and male fitness

There is much indirect evidence from correlational studies that inbreeding reduces male reproductive success [8, 44–46]. In contrast, studies comparing the reproductive output of experimentally created inbred and outbred males have yielded less consistent results. For example, inbreeding depression had no effect on male offspring production in some contexts in flour beetles [*Tribolium castaneum;* 14], while the proportion of offspring sired by inbred males was lower than that of outbred males in bulb mites (*Rhizoglyphus robini*) [*Rhizoglyphus robini*; 15]. In guppies (*Poecilia reticulata*), inbred males sired significantly fewer offspring than outbred males, but only when the inbreeding coefficient was at least *f* =0.25 [i.e. two successive generations of full-sib breeding; 13]. Inbreeding is, in essence, simply a process that decreases heterozygosity. Our experiment therefore reveals a significant heterozygosity-fitness correlation (HFC) for male *G. holbrooki*. However, we also show that detecting this HFC could be difficult using standing variation in heterozygosity, as occurs in field studies [7, 47, 48]. Specifically, we found no effect of residual variation in heterozygosity for either inbred or outbred males. The latter males are roughly equivalent to the field population. It is therefore noteworthy that in a new study of field-caught males, albeit with a larger sample (n = 240 putative sires), we detected a significant HFC for male reproductive success when males compete for females in semi-natural pools (Head et al. *submitted*—available for reviewers in supplementary material). One interpretation of this is that developing under more stressful field conditions exacerbates inbreeding depression.

Studies of inbreeding in wild populations usually fail to tease apart natural and sexual selection against inbred males. Reports of lower reproductive success for less heterozygous (i.e. inbred) males could be due to natural selection because of lower rates of survival [e.g. 49, 50], which will, all else being equal, reduce their lifetime reproductive success; and/or because inbred males are less attractive to females or are weaker competitors [24, 51–53].In our experiment, we can eliminate natural selection through mortality as a source of variation in male reproductive success, so sexual selection most likely explains the lower reproductive success of inbred males. Interestingly, however, in another study we did not detected inbreeding depression for either sperm traits or male attractiveness in *G. holbrooki*, despite much large sample sizes than in the current study (J. Marsh, R. Vega-Trejo, M.L. Head, and M.D. Jennions 'unpublished data'). A lack of inbreeding depression in sperm traits in an introduced species (*G. holbrooki* are feral pests in Australia) could be attributed to low genetic variation due to founder effects [54]. However, the results we present in the current study highlight the need to look how inbreeding affects key fitness components, rather than phenotypic traits that are only indirect proxies for fitness. Given inbreeding depression for male reproductive success, future studies will need to look in more detail at the effects of inbreeding on attractiveness, ejaculate characteristics, male mating behaviour and fertilisation capacity.

### Inbreeding depression in stressful and benign environments

There is a trend for inbreeding depression to be higher in a more stressful environment [25, 26]. By definition a more stressful environment is one that reduces fitness relative to a baseline environment [25]. Our low food diet resulted in almost zero growth over a three-week period [see 36], which strongly suggests that we created a stressful environment. Corroborating this, we have previously shown that this low food diet significantly reduces male attractiveness [55]. In studies of other taxa, mainly insects, a poor juvenile diet has been shown to reduce the ability of males to gain paternity [e.g. 56], which is mainly attributed to a lower sperm count and reduced sperm competitiveness [57, 58]. Elsewhere we have shown that, controlling for age, a poor juvenile diet reduces sperm reserves and sperm replenishment rates in younger male *G. holbrooki* (see Vega-Trejo et al *submitted*— available for reviewers in supplementary material). The males in our current experiment were, however, sufficiently old (28-37 weeks post-maturation) that those on both diets should have had similar sperm production rates so the juvenile diet was not stressful for sperm production. If sperm numbers are a major determinant of male reproductive success this would partly explain why there was no main or interactive effect of diet on male reproductive success. Again, however, this then raises the question of the proximate mechanism causing inbred males to have significantly lower paternity.

Studies of a range of taxa report a weak or no relationship between inbreeding depression and the level of dietary stress [effect size *r* = −0.13 to 0.02; 59, 60, 61], but most of the focal traits measured in these studies are naturally selected. Sexually selected traits that affect male reproductive success are predicted to be more sensitive to inbreeding depression because of their stronger links with fitness [21, 62–64], and more sensitive to environmental stress because they tend to be condition-dependent [65, 66]. It is therefore intriguing that we found significant inbreeding depression for male reproductive success but no effect of diet on a male’s share of paternity. More generally, additional studies of many more taxa are needed to establish whether sexually selected traits show the same pattern as naturally selected traits [25, 26] with respect to whether a more stressful environment elevates inbreeding depression.

### Morphological predictors of male fitness

Males with a relatively long gonopodium for their body size had significantly higher reproductive success, even taking into account the effects of heterozygosity. This corroborates results from another study of *G. holbrooki* in semi-natural pools (Head et al. *submitted*—available for reviewers in supplementary material). Several studies of poeciliid fishes report a link between relative gonopodium length and male fitness [67–71]but see Booksmythe et al. 2016). Male body size is another trait that is often implicated in sexual selection in *G. holbrooki* but in the current study we found that it had no effect on male reproductive success. Male mosquitofish use a coercive mating tactic in which they position themselves behind the female and then thrust their gonopodium forward in an attempt to transfer sperm into the female’s gonoduct [72, 73]. Male size is highly variable and small males have greater manoeuvrability that seems to increase their propensity to sneak copulations [74]. Large males are, however, socially dominant, and might additionally transfer more sperm per encounter because they have larger sperm reserves [75]. This could compensate for the reduced ability of larger males to obtain sneak copulations [74–76]. The net relationship between male body size and reproductive success is likely to depend on the social context, including the absolute size difference between a male and female and the extent of male-male competition for matings [74]. In another paternity study we found that smaller males had significant greater reproductive success when they competed freely for mates in 24 semi-natural pools that varied in the adult sex ratio and habitat complexity (Head et al. *submitted*—available for reviewers in supplementary material). Spatio-temporal variation in how male size affects reproductive success seems likely given the wide size range at which males reach sexual maturity (there is almost no post-maturation growth), even when they are reared under identical laboratory conditions.

## Conclusions

We conducted an experiment that showed that inbreeding reduces a key fitness component (share of paternity) of male *Gambusia holbrooki*. Our design removed most sources of natural selection (e.g. offspring and adult survival, time to maturation), so the lower success of inbred males strongly suggests that inbreeding affects sexual selected traits. This is important as sexual selection against inbred males could reduce the genetic load [33]. If inbred males are less likely to mate and/or fertilize eggs, this will reduce the frequency of deleterious recessive alleles and potentially lower the risk of extinction in small populations [77–80]. This possibility, if generally true in other taxa, could be profitably incorporated into models of population viability.Our study is also a reminder that standing variation in heterozygosity plays an important role in the likelihood of detecting inbreeding depression, which might explain variation in reported level of inbreeding depression in other studies [e.g. 7, 47, 48]. Standing variation in heterozygosity, hence the use of heterozgosity-fitness correlations, was insufficient to detect inbreeding depression in our study as there was no effect of relative heterozygosity on paternity. We only detected inbreeding depression between our inbreeding treatment led to a 23% decline in heterozygosity. Given the potential for inbreeding to shape the evolution of key life history traits [81], more studies are needed that quantify inbreeding depression by taking an experimental approach and then measure fitness as directly as possible (i.e. reproductive success not simply phenotypic traits).

### Authors' contributions

R.V.T., M.L.H., and M.D.J. designed the study. R.V.T carried out the experimental work. J.S.K. analysed the paternity data. R.V.T., M.L.H., and M.D.J. analysed the data and wrote the manuscript. All the authors contributed substantially to revisions, and gave final approval for publication.

## Acknowledgements

We thank the ANU Animal Services team for fish maintenance. Animal use permit: ANU AEEC animal ethics protocol A2011/64. The authors thank Rose E. O’Dea for helping with the experimental work.

